# Novel Analgesic Agents Obtained by Molecular Hybridization of Orthosteric and Allosteric Ligands

**DOI:** 10.1101/866012

**Authors:** Carlo Matera, Lisa Flammini, Fabio Riefolo, Giuseppe Domenichini, Marco De Amici, Elisabetta Barocelli, Clelia Dallanoce, Simona Bertoni

## Abstract

Despite the high incidence of acute and chronic pain in the general population, the efficacy of currently available medications is unsatisfactory. Insufficient management of pain has a profound impact on the quality of life and can have serious physical, psychological, social, and economic consequences. This unmet need reflects a failure to develop novel classes of analgesic drugs with superior clinical properties and lower risk of abuse. Nevertheless, recent advances in our understanding of the neurobiology of pain are offering new opportunities for developing different therapeutic approaches. Among those, the activation of M2 muscarinic acetylcholine receptors, which play a key role in the cholinergic regulation of the nociceptive transmission, constitutes one of the most promising strategies. We have recently developed a small library of novel pharmacological agents by merging the structures of known orthosteric and allosteric muscarinic ligands through their molecular hybridization, an emerging approach in medicinal chemistry based on the combination of pharmacophoric moieties of different bioactive substances to produce a new compound with improved pharmacological properties. Herein we report the functional characterization of the new ligands in vitro and the assessment of their efficacy as analgesic agents and tolerability in mice. This work provides new insights for the development and optimization of novel muscarinic hybrid compounds for the management of pain.

## INTRODUCTION

Pain, which is defined as “an unpleasant sensory and emotional experience that emerges as a normal response to injury and is a symptom of many diseases”, is the most common reason people seek medical care.(Fiorino and Garcia-Guzman, 2012) The efficacy of currently available medications is unsatisfactory owing to their limited effect size and the low responder rate. Insufficient management of pain has a profound impact on the quality of life and can have serious physical, psychological, social, and economic consequences.(King and Fraser, 2013) However, progress by the pharmaceutical industry in exploiting new mechanisms for clinical efficacy has been limited, and pain treatments have changed little since the introduction of nonsteroidal anti-inflammatory agents more than three decades ago, especially because of our incomplete understanding of the pathophysiology of human pain.(Fiorino and Garcia-Guzman, 2012) Thus, there is a large unmet need for innovative therapies for this condition, especially in case of chronic pain.

Within the rather complex network of pathways that contribute to the modulation of pain transmission, G protein-coupled receptors (GPCRs) exert a relevant role. Indeed, activation of different GPCRs, including opioid, cannabinoid, α2-adrenergic, muscarinic, *γ*-aminobutyric acid (GABAB), metabotropic glutamate, and somatostatin receptor families, may produce analgesic effects.(Pan et al., 2008) The activation of muscarinic acetylcholine receptors (mAChRs) constitutes one of the most promising strategies to develop novel analgesic agents.(De Angelis and Tata, 2016; Eisenach, 1999; Jones and Dunlop, 2007; Matera and Tata, 2014; Tata, 2008) These receptors are classified into five distinct subtypes, denoted as M1– M5. Three of these receptor subtypes (M1, M3, M5) have been shown to couple to G proteins of the G_q/11_ family, whereas the remaining two subtypes (M2, M4) preferentially signal through the G_i/o_ family of G proteins.(Kruse et al., 2014) Previous studies have provided strong evidence that M2 mAChRs play a key role in the cholinergic regulation of nociceptive transmission. It has been found that the M2 subtype is the main muscarinic receptor subtype in the spinal cord dorsal horn,(Pan et al., 2008) and that also dorsal root ganglia (DRG) neurons express high levels of M2 mRNA. The upregulation of the M2 subtype in DRG neurons has also been reported in rats with neuropathic pain induced by nerve injury.(Clayton et al., 2007; De Angelis and Tata, 2016; De Angelis et al., 2014; Hayashida et al., 2006)

In the framework of a research project focused on the development of unprecedented muscarinic ligands, we had previously prepared and studied a new series of hybrid agonists. Molecular hybridization is an emerging concept in drug design based on the combination of pharmacophoric moieties of different bioactive substances to generate a new hybrid compound with an enhanced pharmacological profile, when compared to the parent drugs, in terms of affinity, efficacy, selectivity or safety/tolerability.(Decker, 2017; Matera et al., 2019; 2015; Mohr et al., 2010; Viegas-Junior et al., 2007) Our muscarinic hybrid compounds were obtained by combining in the same molecular structure two distinct pharmacophore elements belonging to (a) potent orthosteric agonists and (b) subtype-selective allosteric ligands, in order to investigate how the introduction of such allosteric moieties could affect the binding topology of the orthosteric agonists (Figure 1).(De Amici et al., 2010; Disingrini et al., 2006) We found that these hybrid compounds are characterized by a truly bitopic (orthosteric and allosteric) binding mode (hence the term “dualsteric” ligands),(Antony et al., 2009) as well as other distinctive features when compared to their parent compounds, such as receptor subtype selectivity and biased agonism.(Bock et al., 2016; 2012; De Min et al., 2017) We also demonstrated that some of these hybrid compounds display interesting pharmacological properties,(Cristofaro et al., 2018; Riefolo et al., 2019) including remarkable analgesic effects in vivo in mice. Noteworthy, one of these compounds, from now on named Iper-8-N (Figure 1, compound *8b* in the reporting article), even combined the potent antinociceptive activity with the absence of relevant cholinergic side effects.(Matera et al., 2014)

**Figure 1.**
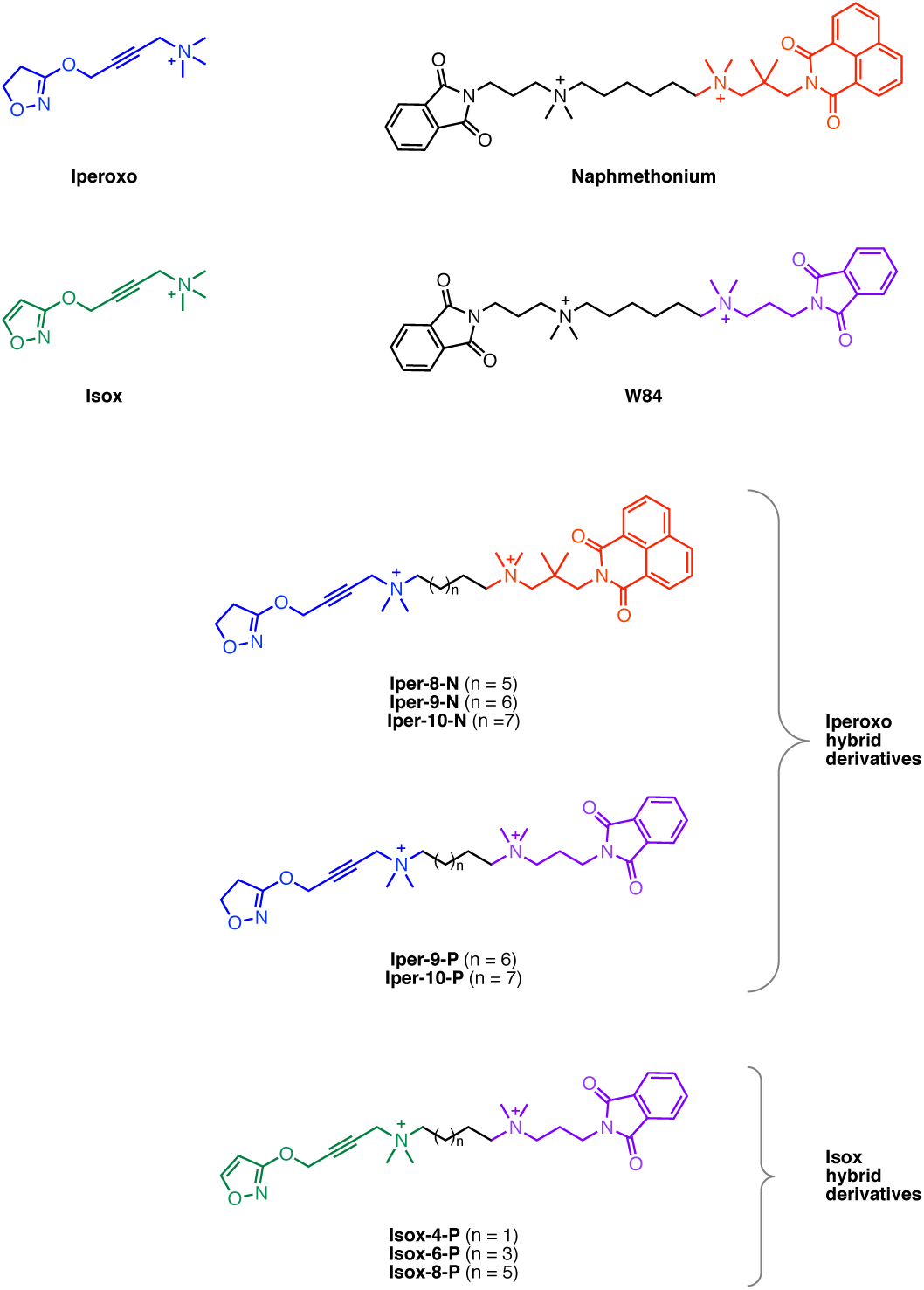
Chemical structures of the muscarinic ligands discussed in this work: the orthosteric agonists Iperoxo and Isox, the allosteric modulators Naphmethonium and W84, and their corresponding hybrid compounds. “N” and “P” stand for naphthalimide and phtalimide (from Naphmethonium and W84), respectively.

We have recently expanded our toolbox of hybrid muscarinic ligands and performed a preliminary assessment of their pharmacological profile in vitro.(Messerer et al., 2017) Among those, two groups of molecules were selected for further investigation. The new sets of compounds are distinguished from others previously described(Matera et al., 2014) for (a) the length of the polymethylene spacer connecting the orthosteric and the allosteric moieties (Iperoxo derivatives, Figure 1), or (b) the nature of the orthosteric moiety (Isox derivatives, Figure 1). We report here their functional characterization on isolated tissue preparations, the evaluation of their antinociceptive activity in vivo as well as the quantification of their potential muscarinic-mediated side effects. This work further expands the knowledge of the chemical space around hybrid muscarinic ligands and should facilitate the design of novel pharmacological agents for the management of pain.

## RESULTS AND DISCUSSION

All the compounds used in our experiments were prepared as previously reported.(Messerer et al., 2017) We initially evaluated the ability of the new derivatives to activate the target receptor M2 with experiments in isolated tissue preparations of guinea pig left atrium (Table 1). The four Iperoxo-containing ligands (Iper-9-N, Iper-10-N, Iper-9-P, Iper-10-P) behaved as potent M2 mAChR full agonists, although they resulted from 1.5 to 2 orders of magnitude less potent than their corresponding orthosteric agonist Iperoxo. Regarding the Isox derivatives, we observed a complete loss of agonist activity at the M2 mAChR for Isox-4-P and Isox-8-P, while a positive inotropic effect appeared for all three compounds, in stark contrast with the pharmacological profile of their corresponding orthosteric agonist Isox. Only Isox-6-P showed a very weak negative inotropic effect up to 10 μM.

**Table 1.** In vitro functional activity at M2 muscarinic receptors (guinea pig left atrium) of the muscarinic hybrid compounds and the corresponding orthosteric agonists described in this work.

| compound | pEC<sub>50</sub><sup>a</sup> | i.a.<sup>b</sup> | pK<sub>B</sub><sup>c</sup> |
| --- | --- | --- | --- |
| Iperoxo | 10.10 (0.13)<sup>d</sup> | 1.00<sup>d</sup> | - |
| Iper-8-N | 8.15 (0.09)<sup>e</sup> | 1.01<sup>e</sup> | - |
| Iper-9-N | 8.67 (0.09) | 1.18 | - |
| Iper-10-N | 8.40 (0.09) | 1.07 | - |
| Iper-9-P | 8.28 (0.06) | 0.91 | - |
| Iper-10-P | 8.74 (0.03) | 1.11 | - |
| Isox | 8.54 (0.03)<sup>d</sup> | 1.00<sup>d</sup> | - |
| Isox-4-P | positive inotropic<sup>f</sup> | - | inactive<sup>g</sup> |
| Isox-6-P | positive inotropic<sup>h</sup> | 0.45 | inactive<sup>g</sup> |
| Isox-8-P | positive inotropic<sup>i</sup> | - | 6.41 (0.01) |

Data are expressed as mean (SEM) of 2–5 experiments. <sup>a</sup>pEC<sub>50</sub> (SEM) values are the negative logarithm of the agonist concentration that caused 50% of the maximum response. <sup>b</sup>i.a. = intrinsic activity ( $\alpha$ ) measured as the ratio between the maximum negative inotropic effect of the compound and the maximum response of the reference full agonist bethanechol. <sup>c</sup>Apparent pK<sub>B</sub> values (SEM) were calculated according to (Furchgott and Bursztyn, 1967) <sup>d</sup>Data taken from (Dallanocce et al., 1999) <sup>e</sup>Data taken from (Matera et al., 2014) <sup>f</sup>Starting from 1 μM. <sup>g</sup>Up to 10 μM. <sup>h</sup>Starting from 30 μM. <sup>i</sup>Starting from 100 nM.

Overall, when compared to the best performing compound of the previous series (Iper-8-N),(Matera et al., 2014) the four novel Iperoxo-like hybrids maintained a remarkable efficacy and potency, regardless the nature of the allosteric portion and the length of the linker, whereas the three Isox derivatives nearly completely lost efficacy at M2.

The two sets of compounds under study were then assayed in mice for their antinociceptive effect by means of the writhing test, a conventional chemical method used to induce pain of peripheral origin by intraperitoneal injection of 0.6% acetic acid solution (chemical algesia),(Koster et al., 1959) and the hot plate test, a method that is generally used for testing the effectiveness of centrally acting analgesic by observing the reaction time to pain caused by heat (thermal algesia).(Eddy and Leimbach, 1953)

All Iperoxo derivatives showed a potent and dose-dependent analgesic activity in the writhing test, except for compound Iper-9-P, which emerged as the less potent antinociceptive agent of the group (Figure 2). In fact, Iper-9-P did not show analgesic effects up to a dose of 0.1 mg/kg and exhibited only a mild analgesic activity (56%) at 1 mg/kg. Conversely, the hybrids Iper-10-P, Iper-9-N and Iper-10-N displayed a remarkable and significant analgesic activity at 0.1 mg/kg, inhibiting by 86% (ID_50_ = 22.1 μg/kg), 76% (ID_50_ = 25.8 μg/kg) and 84% (ID_50_ = 7.0 μg/kg), respectively, the acetic acid-induced writhing response, similarly to the reference hybrid agonist Iper-8-N, which had given an 80% inhibition of the writhing response at 0.5 mg/kg.(Matera et al., 2014)

**Figure 2.**
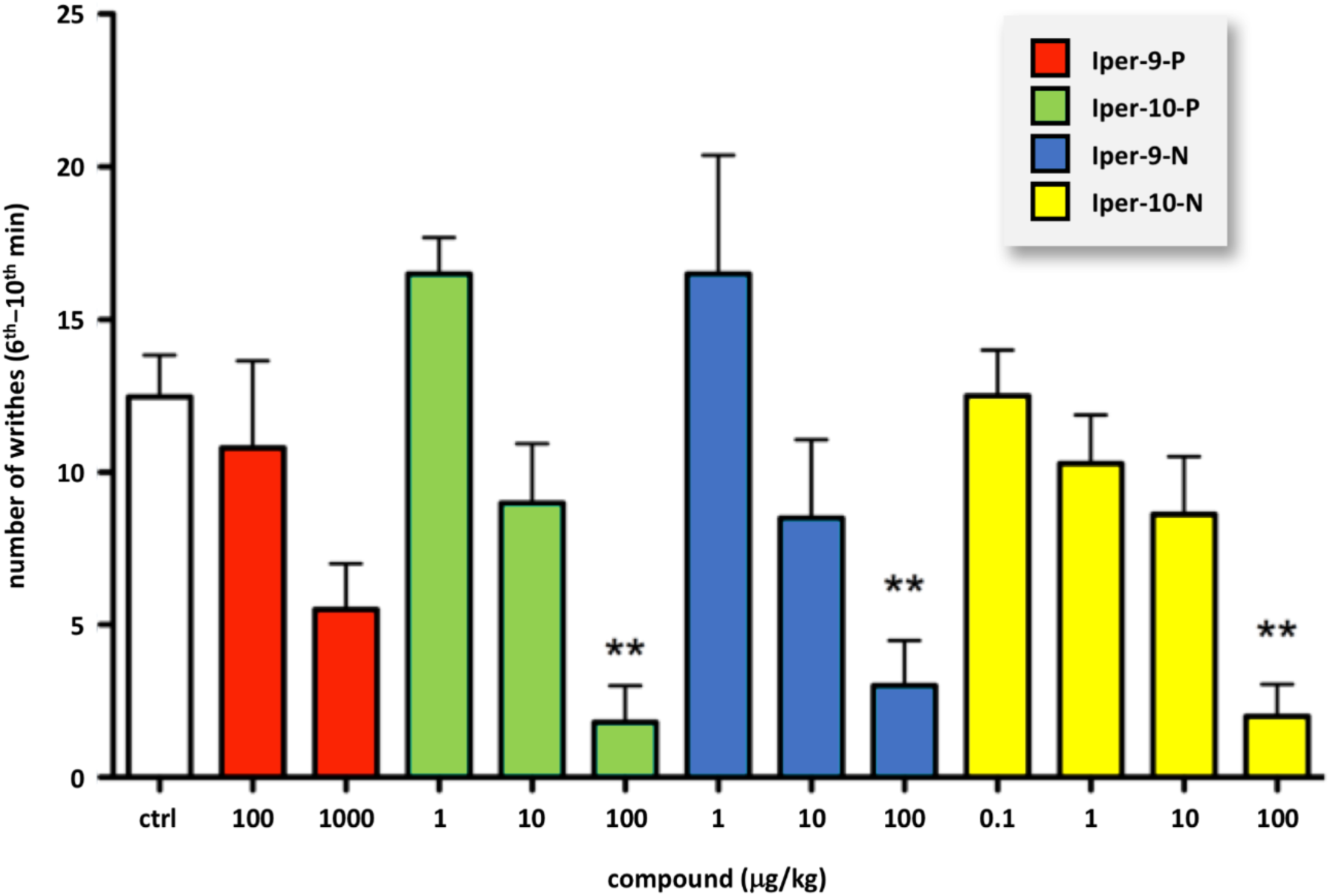
Writhing test with Iperoxo hybrids. Number of acetic acid-induced writhes observed (6^th^–10^th^ min) in mice subcutaneously administered with vehicle (control group, ctrl) or with Iperoxo hybrid derivatives (μg/kg). Data are expressed as mean ± SEM of 4–8 animals and were analyzed by one-way ANOVA followed by Dunnett’s post-hoc test for statistical significance, (**) p < 0.01 vs control group.

The most promising compounds (Iper-10-P, Iper-9-N, Iper-10-N) were then tested in the hot plate test at 0.1 mg/kg, i.e. the dose that gave the maximum analgesic effect in the writhing test. Interestingly, only Iper-10-P gave a significant increase in the time of latency in comparison with control, delaying the response of about 10 s at 30 min after administration of the drug. On the other hand, none of the two Naphmethonium-related derivatives, Iper-9-N and Iper-10-N, was able to modify the response latency to the thermal stimulus (Figure 3).

**Figure 3.**
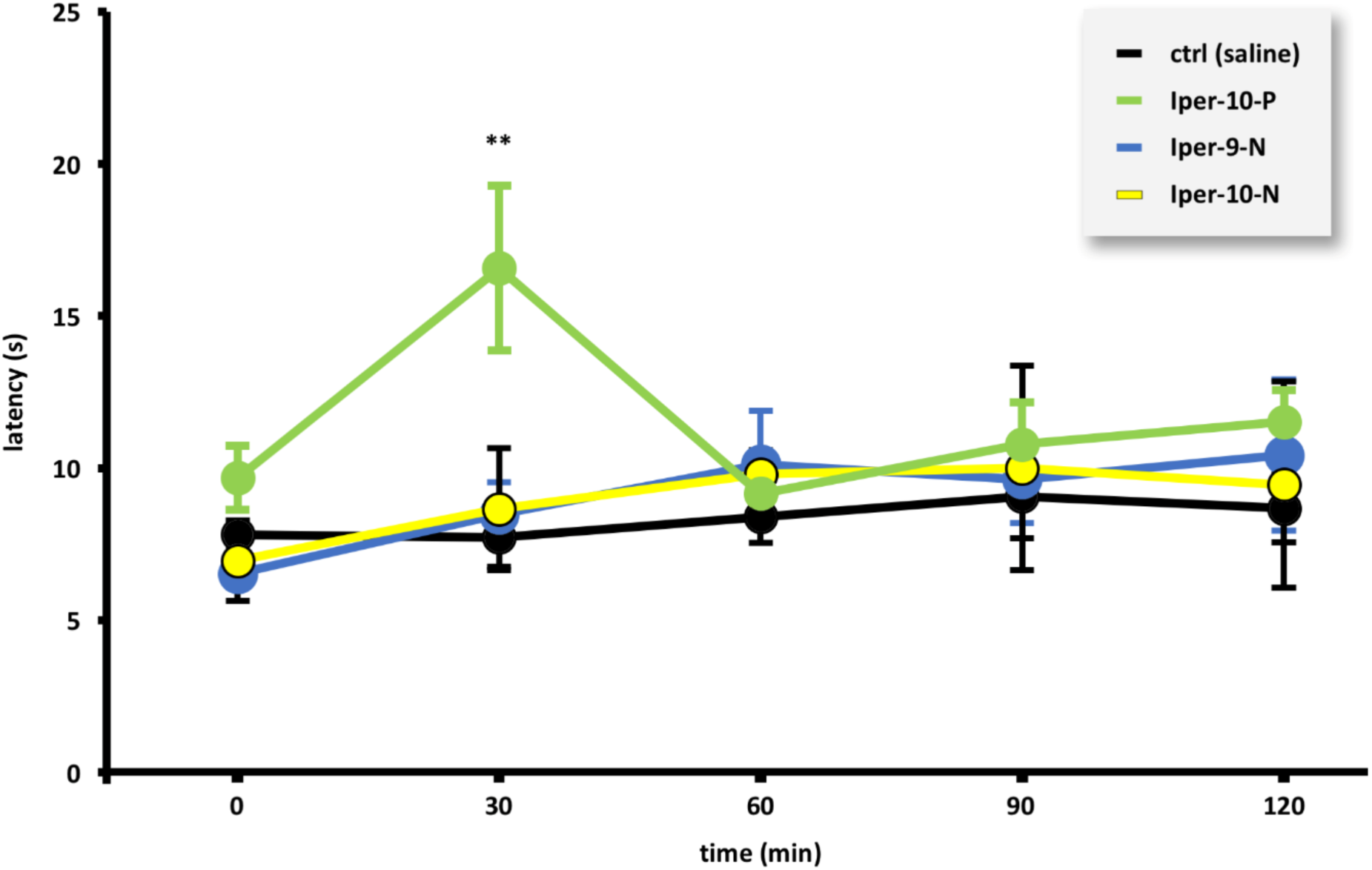
Hot plate test with Iperoxo hybrids. Thermal response latency in mice subcutaneously administered with vehicle (control group, ctrl) or with Iperoxo hybrid derivatives Iper-10-P, Iper-9-N, and Iper-10-N (100 μg/kg). Analgesia was quantified as hot-plate latency (s) over the time (min). Data are expressed as mean ± SEM of 4–6 observations and were analyzed by two-way ANOVA followed by Bonferroni’s post-hoc test for statistical significance, (**) p < 0.01 vs control group.

Since potential sedative effects could interfere with the data collected in the nociception assays, these three Iperoxo hybrids were further examined in the open field test in mice, an established method to assay general locomotor activity levels, anxiety, and willingness to explore of the animals.(Gould et al., 2009)

Among the tested compounds, only Iper-10-P produced a decrease of the spontaneous motility of the animals at 100 μg/kg, while no effects were produced by the other derivatives, either on the locomotor activity or on the duration of immobility time (Table 2), highlighting Iper-10-N as the best performing compound.

**Table 2.** Locomotor activity and immobility time in mice subcutaneously administered with vehicle (ctrl) or with Iperoxo hybrid derivatives Iper-10-P, Iper-9-N, and Iper-10-N.

| compound ($\mu\text{g/kg}$) | locomotor activity (m) | immobility time (min) |
| --- | --- | --- |
| ctrl | $40.9 \pm 1.9$ | $39.2 \pm 1.1$ |
| Iper-10-P (10) | $48.3 \pm 5.6$ | $35.1 \pm 2.0$ |
| Iper-10-P (50) | $42.8 \pm 4.7$ | $37.0 \pm 2.4$ |
| Iper-10-P (100) | $22.5 \pm 2.7^*$ | $45.0 \pm 2.0$ |
| Iper-9-N (10) | $54.8 \pm 13.6$ | $32.2 \pm 7.4$ |
| Iper-9-N (50) | $33.4 \pm 3.8$ | $41.4 \pm 2.4$ |
| Iper-9-N (100) | $27.8 \pm 3.7$ | $42.1 \pm 2.4$ |
| Iper-10-N (10) | $60.6 \pm 3.7$ | $33.0 \pm 0.8$ |
| Iper-10-N (50) | $46.8 \pm 5.1$ | $36.1 \pm 2.0$ |
| Iper-10-N (100) | $35.8 \pm 2.5$ | $39.8 \pm 1.0$ |

Data are expressed as mean ± SEM of 3–10 observations and were analyzed by one-way ANOVA followed by Bonferroni's post-hoc test for statistical significance, (\*) $p < 0.05$ vs control group.

As for the Isox hybrids, all compounds under study showed a dose-dependent analgesic activity in the writhing test, although they resulted much less potent in comparison with both their parent orthosteric agonist Isox(Barocelli et al., 2001) and the Iperoxo-like hybrids (Figure 4). Interestingly, we observed that the capacity of the Isox hybrids to prevent nociception was correlated to the length of the polymethylene spacer, which is in agreement with our previous findings on similar analogs.(Matera et al., 2014) In fact, Isox-4-P, the shortest ligand of this set, was the compound with the lowest antinociceptive activity, displaying significant analgesic effects (82% inhibition) only at the dose of 40 mg/kg (ID_50_ = 26.6mg/kg). On the other hand, the two homologs characterized by longer linkers, Isox-6-P and Isox-8-P, showed remarkable analgesic activity already at 10 mg/kg, inhibiting by 60% (ID_50_ = 8.4mg/kg) and 90% (ID_50_ = 6.6mg/kg), respectively, the acetic acid-induced writhing response at that dose.

**Figure 4.**
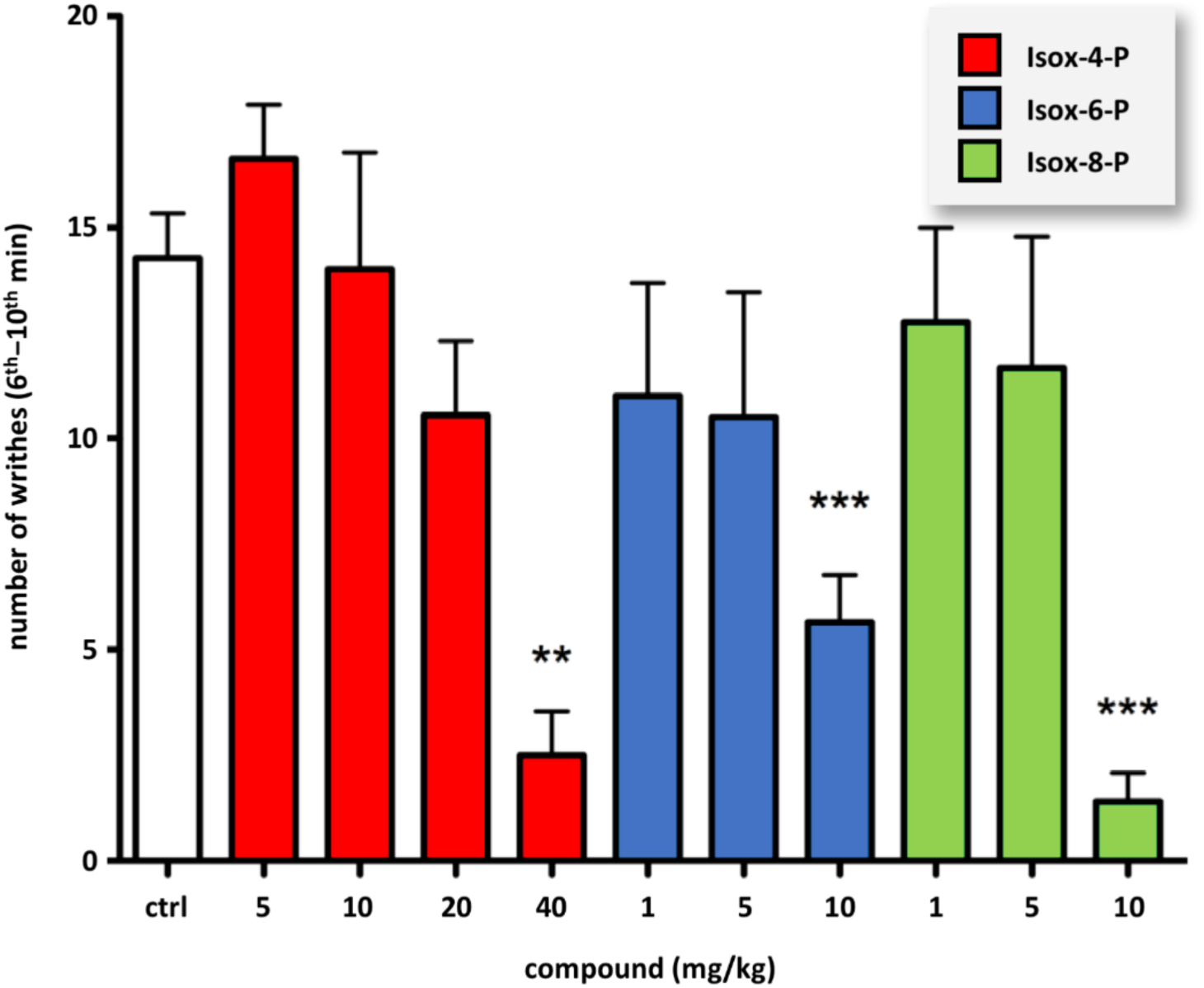
Writhing test with Isox hybrids. Number of acetic acid-induced writhes observed (6^th^–10^th^ min) in mice subcutaneously administered with vehicle (control group, ctrl) or with Isox hybrid derivatives (mg/kg). Data are expressed as mean ± SEM of 4–8 animals and were analyzed by one-way ANOVA followed by Dunnett’s post-hoc test for statistical significance, (**) p < 0.01 vs control group, (***) p < 0.001 vs control group.

We next studied the antinociceptive activity of the Isox hybrids also in the hot plate test at the dose of 10 mg/kg (Figure 5). None of the hybrid ligands produced an increase of the hot plate latency times in comparison with the control, thus resulting devoid of analgesic effects in this type of assay.

**Figure 5.**
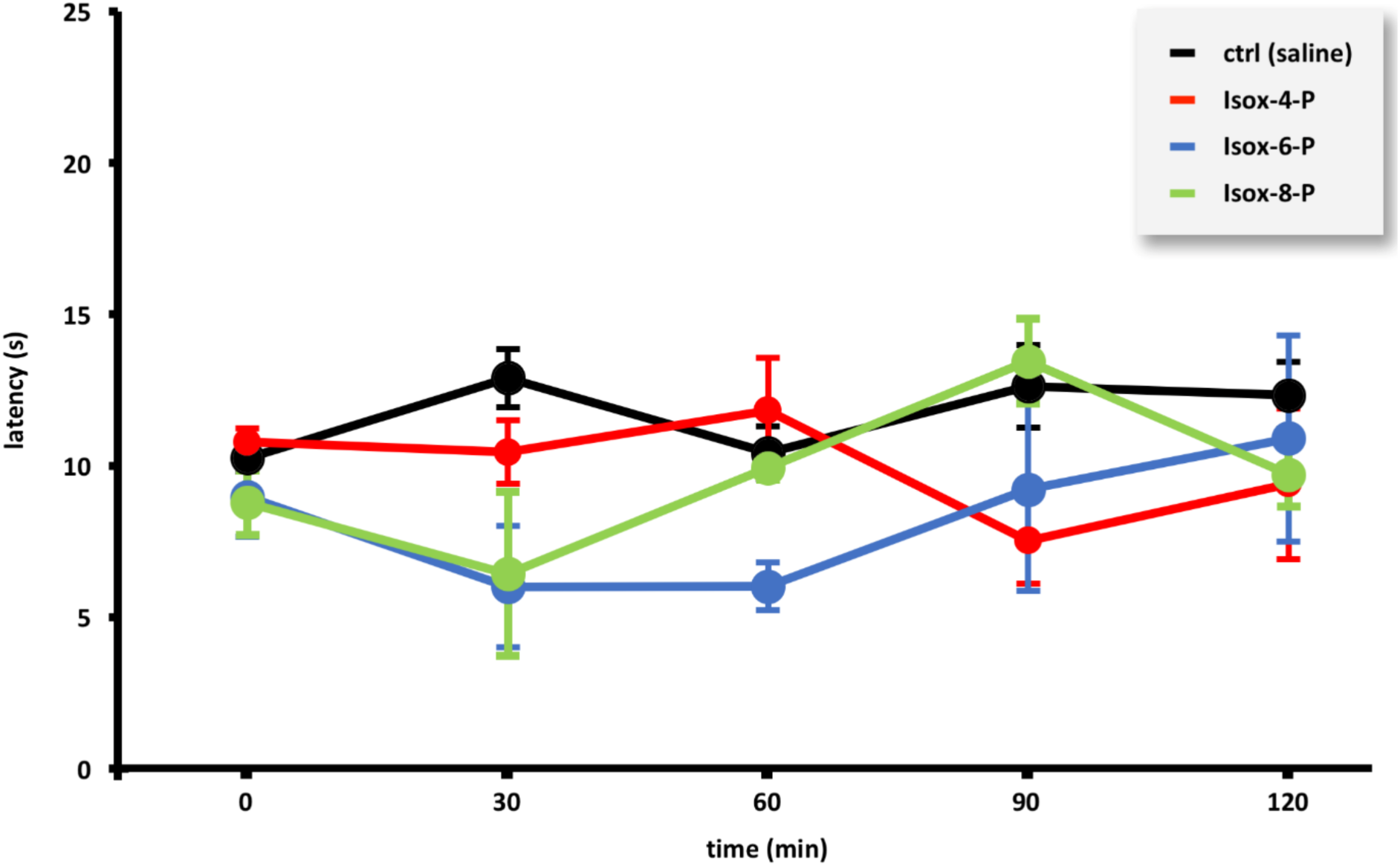
Hot plate test with Isox hybrids. Thermal response latency in mice subcutaneously administered with vehicle (control group, ctrl) or with the Isox hybrid derivatives Isox-4-P, Isox-6-P, Isox-8-P (10 mg/kg). Analgesia was quantified as hot-plate latency (s) over the time (min). Data are expressed as mean ± SEM of 4–6 observations and were analyzed by two-way ANOVA followed by Bonferroni’s post-hoc test for statistical significance, (**) p < 0.01 vs control group.

In order to identify potential false positives also in this set of compounds, we examined the Isox derivatives in the open field test (data not shown). Among the derivatives under study, only Isox-4-P showed significant sedative effects, as denoted by both the decrease of the distance travelled (23.1 ± 5.4 m vs 53.3 ± 3.3 m for the control group, p < 0.05) and the increase of the immobility time at the dose of 40 mg/kg (47.5 ± 2.0 vs 32.0 ± 3.0 for the control group, p < 0.05), while Isox-8-P emerged as the most promising compound of the group, as it combined an effective analgesic activity with the absence of sedative effects.

Finally, to evaluate the tolerability profile of the compounds under investigation, we estimated the incidence of major muscarinic side effects such as diarrhea, salivation, lachrymation, hypothermia and tremor following their subcutaneous administration at the doses that gave analgesic effects in mice (Table 3). All the novel Iperoxo hybrids (Iper-9-N, Iper-10-N, Iper-9-P, Iper-10-P) displayed generally a considerable presence of unwanted muscarinic side effects. In comparison with the best performing agent from the previous work, Iper-8-N,(Matera et al., 2014) the elongation of the polymethylene spacer from 8 to 9 or 10 carbon atoms afforded compounds endowed with greater analgesic potency but also a much worse tolerability profile, regardless the nature of the allosteric moiety. On the contrary, the Isox hybrids (Isox-4-P, Isox-6-P, Isox-8-P) were all characterized by a good profile of tolerability. In fact, unlike their parent orthosteric agonist Isox,(Barocelli et al., 2001) these compounds resulted all devoid of muscarinic adverse effects, except for the hypothermia, which was detected in about 50% of the animals treated with Isox-4-P and Isox-6-P, and in all the animals treated with Isox-8-P.

**Table 3.** Unwanted muscarinic side effects produced by subcutaneous administration of the muscarinic hybrid compounds and their corresponding parent orthosteric agonists at doses that significantly inhibited acetic acid-induced writhing responses.

| compound | dose (mg/kg) | diarrhea | salivation | lachrymation | hypothermia | tremor |
| --- | --- | --- | --- | --- | --- | --- |
| Iperoxo<sup>a</sup> | 0.001 | - | + | - | ++ | - |
| Iper-8-N<sup>a</sup> | 0.01 | - | - | - | - | - |
| Iper-9-N | 1 | ++++ | ++ | ++ | ++++ | - |
| Iper-10-N | 0.1 | ++++ | ++++ | ++++ | +++ | - |
| Iper-9-P | 0.1 | ++ | ++ | + | +++ | - |
| Iper-10-P | 0.1 | ++ | +++ | + | ++++ | - |
| Isox-4-P | 40 | - | - | - | ++ | - |
| Isox-6-P | 10 | - | - | - | ++ | - |
| Isox-8-P | 10 | - | - | - | ++++ | - |

<sup>a</sup>Data taken from (Matera et al., 2014) (++++), (+++), (++) and (+) indicate 100%, 75%, 50%, and 25% incidence of the side effect, respectively. (-) indicates absence of the side effect.

It is worth noting that none of the compounds under investigation in both sets showed tremorigenic effects, which supports the idea that such bis-quaternary ammonium agents are characterized by a purely peripheral mechanism of action, and hence that their analgesic (and hypothermic) effects may originate peripherally rather than centrally.(Sánchez and Meier, 1993)

## CONCLUSION

In a previous study, we had found that analgesic agents designed by hybridization of potent orthosteric agonists and subtype-selective allosteric modulators of M2 muscarinic receptors are generally characterized by a promising therapeutic profile in terms of efficacy and tolerability. Here, in the attempt to further explore the chemical space around this type of muscarinic ligands and identify candidates characterized by enhanced properties, we studied the analgesic activity of two novel sets of hybrid compounds as well as their tolerability profile in vivo.

All the new ligands obtained by hybridization of Iperoxo behaved as potent full agonists at the M2 mAChR in vitro, with comparable level of potency and efficacy throughout the series, whereas they gave different outcomes when assayed as antinociceptive agents in mice. In particular, Iper-10-P, Iper-9-N and Iper-10-N emerged as the most potent members of the set in the writhing test, while only Iper-10-P resulted active in the hot plate test. While the sedative effects detected in the open field test for Iper-10-P could cast some doubts on its analgesic action, we believe that Iper-10-N can be considered the most interesting compound of this set, since it is characterized by both high analgesic potency (ID_50_ = 7.0 μg/kg) and no sedative effects in mice. Unfortunately, it did not show an equally good tolerability profile at the active dose, unlike its predecessor Iper-8-N. Thus, we can conclude that the further elongation of the polymethylene spacer connecting the orthosteric and allosteric moieties (from 8 to 9 and 10 carbon atoms) with respect to the reference compound Iper-8-N is not advantageous, since it only moderately improves the analgesic potency at the expense of the tolerability. This could mean that an optimal length may have been already identified for these model ligands and suggests to modify only the nature of the spacer as the next step to improve their putative usefulness as analgesic agents.

On the other hand, the three compounds obtained by hybridization of Isox (Isox-4-P, Isox-6-P, Isox-8-P) showed interesting analgesic properties in the writhing test in mice, especially Isox-8-P, together with a remarkably good profile of tolerability, although much higher doses were needed to observe significant antinociceptive effects in comparison with their parent orthosteric agonist and with the Iperoxo hybrids. Intriguingly, all the Isox hybrids were unable to activate M2 mAChRs in our functional assays in vitro, which suggests that the mechanism at the basis of their analgesic effects could be purely allosteric in nature or associated to a different receptor subtype or family. Further studies will be required to identify the true mechanism of action of such compounds.

## MATERIALS AND METHODS

### Animals

*In vitro* assays were conducted on electrically-stimulated left atria from male albino guinea pigs (250–350 g; Charles River, Italy). Behavioral experiments were conducted on male Swiss mice (25–35 g; Charles River, Italy). Animals were housed, handled and cared for in accordance with the European Community Council Directive 86 (609) EEC, and the experimental protocols were carried out in compliance with Italian regulations (DL26/2014) and with the local Ethical Committee Guidelines for Animal Research. All procedures were carried out from 9 am to 2 pm in animals fasted 18 h before each experiment.

### Functional studies on guinea pig left atrium

Isolated preparations were set up following the techniques previously described.(Barocelli et al., 2000) Heart was rapidly dissected and the right and left atria separated. The left atrium was suspended under 0.5 g tension in a 20-mL organ bath in a modified Krebs-Henseleit buffer solution (mM composition: NaCl 118.9, KCl 4.6, CaCl_2_ 2.5, KH_2_PO_4_ 1.2, NaHCO_3_ 25, MgSO_4_·7H_2_O 1.2, glucose 11.1) gassed with carbogen (95% O_2_ ± 5% CO_2_) at 32 °C. The tissue was electrically-paced by rectangular submaximal impulses (2 Hz, 5 ms, 5 V) by using platinum wire electrodes. After a 30-min stabilization period, inotropic responses were measured as changes in isometric tension. Agonist concentration–response curves were constructed in each tissue by cumulative application of concentrations of the test compounds (1 nM–30 μM). The agonist potency was expressed as pEC_50_ (-logEC_50_) calculated by linear regression analysis using the least square method. Intrinsic activity (α) was calculated as a fraction of the maximal response to the reference full agonist bethanechol. Concentration-response curves of the agonists were reconstructed in the presence of atropine 0.1 μM. When the compounds were tested as antagonists, a dose– response curve to the full agonists bethanechol was repeated after 30 min incubation with the test compounds (1–30 μM). The potency of the compounds acting as antagonists is expressed as p*K*_B_ value, the calculated molar concentration of the test compounds that causes a twofold increase in the EC_50_ values of bethanechol, calculated according to Furchgott’s method.(Furchgott and Bursztyn, 1967)

### Writhing test

The writhing test was performed according to Koster et al.(Koster et al., 1959) Briefly, 30 min after the compounds under study or the vehicle (saline 0.9%, 10 mL/kg) were subcutaneously (s.c.) administered, groups of 6–8 mice were intraperitoneally (i.p.) injected with 0.6% acetic acid and placed in observational chambers. The number of writhes of each mouse was counted in a period of 6–10 min after the application of the algogen agent. For each compound under study, the dose able to reduce the number of acetic acid-induced writhes by 50% in comparison with those of the control group, exposed to the algogen agent but treated with vehicle, was calculated and expressed as ID_50_.

### Hot plate test

The hot plate test was performed according to the method described by Woolfe and Macdonald.(Woolfe and MacDonald, 1944) Mice were individually placed on the 55 °C hot plate apparatus [Model 475, Technical Lab Instruments Inc, Pequannock (NJ), USA] and the time between the placement and the occurrence of anterior paw licking, shaking or jumping was recorded as “latency time” (s). A 30-s cut-off time was set to prevent tissue damage. In order to exclude hypo- or hyper-sensitive mice, animals with latency time shorter than 10 s or longer than 18 s were eliminated from the study. Mice received subcutaneously the compounds under study or the vehicle 30 min before the assay: the latency time was measured 30, 60, 90 and 120 min after the compounds or the vehicle injection.

### Open field test

Spontaneous locomotor activity was assessed in the open field apparatus, consisting in four square chambers (height 40 cm, width 45 cm, depth 45 cm) with a video camera fixed above them. Each animal, 30 min before the trial, received subcutaneously the compounds under study or the vehicle. Locomotor activity was recorded for 60 min by means of a video-tracking software (Any-maze, Ugo Basile, Comerio, Italy) measuring the total traveled distance (m) and the total time of immobility (min).

### Cholinergic side effects

The incidence of cholinergic unwanted side effects was simultaneously estimated in mice during the writhing test. Body temperature was recorded with a rectal probe (Delta Ohm HD8704, Padova, Italy) immediately before and 30 min after drug administration in an air-conditioned room (20 °C); the hypothermic effect was scored as present if body temperature decreased by 2 °C with respect to basal temperature. The presence or absence of sialagogic, lachrymatory and tremorigenic responses and of intestinal hypermotility were recorded on a period of 1 h following s.c. administration of the test compounds. Salivation and lachrymation were scored as present if the area surrounding the mouth or the eyes were wet, while tremor was scored as present on the basis of the tremor provoked by handling during the temperature measurement. Results are expressed as percentage of treated animals exhibiting the specific cholinergic side effect.

### Data analysis

All data are given as mean ± SEM. ID_50_ values of analgesic activity (i.e., the dose that inhibits acetic acid writhing response by 50% compared to the control group) were calculated by non-linear regression analysis (Prism 5.0, GraphPad Software, San Diego, CA, USA). Statistical analysis was performed as indicated in the captions. A p value < 0.05 was considered significant, a p value < 0.01 was considered highly significant.

## ACKNOWLEDGMENTS

This research received no specific grant from any funding agency.

## REFERENCES

Antony, J., Kellershohn, K., Mohr-Andrä, M., Kebig, A., Prilla, S., Muth, M., Heller, E., Disingrini, T., Dallanoce, C., Bertoni, S., Schrobang, J., Tränkle, C., Kostenis, E., Christopoulos, A., Höltje, H.-D., Barocelli, E., De Amici, M., Holzgrabe, U., Mohr, K., 2009. Dualsteric GPCR targeting: a novel route to binding and signaling pathway selectivity. FASEB J. 23, 442–450. doi:10.1096/fj.08-114751

Barocelli, E., Ballabeni, V., Bertoni, S., Dallanoce, C., De Amici, M., De Micheli, C., Impicciatore, M., 2000. New analogues of oxotremorine and oxotremorine-M: estimation of their in vitro affinity and efficacy at muscarinic receptor subtypes. Life Sci. 67, 717–723. doi:10.1016/s0024-3205(00)00661-5

Barocelli, E., Ballabeni, V., Bertoni, S., De Amici, M., Impicciatore, M., 2001. Evidence for specific analgesic activity of a muscarinic agonist selected among a new series of acetylenic derivatives. Life Sci. 68, 1775–1785. doi:10.1016/s0024-3205(01)00973-0

Bock, A., Bermudez, M., Krebs, F., Matera, C., Chirinda, B., Sydow, D., Dallanoce, C., Holzgrabe, U., De Amici, M., Lohse, M.J., Wolber, G., Mohr, K., 2016. Ligand binding ensembles determine graded agonist efficacies at a G protein-coupled receptor. Journal of Biological Chemistry 291, 16375–16389. doi:10.1074/jbc.M116.735431

Bock, A., Merten, N., Schrage, R., Dallanoce, C., Bätz, J., Klöckner, J., Schmitz, J., Matera, C., Simon, K., Kebig, A., Peters, L., Müller, A., Schrobang-Ley, J., Tränkle, C., Hoffmann, C., De Amici, M., Holzgrabe, U., Kostenis, E., Mohr, K., 2012. The allosteric vestibule of a seven transmembrane helical receptor controls G-protein coupling. Nature Communications 3. doi:10.1038/ncomms2028

Clayton, B.A., Hayashida, K.-I., Childers, S.R., Xiao, R., Eisenach, J.C., 2007. Oral donepezil reduces hypersensitivity after nerve injury by a spinal muscarinic receptor mechanism. Anesthesiology 106, 1019–1025. doi:10.1097/01.anes.0000265163.22007.6d

Cristofaro, I., Spinello, Z., Matera, C., Fiore, M., Conti, L., De Amici, M., Dallanoce, C., Tata, A.M., 2018. Activation of M2 muscarinic acetylcholine receptors by a hybrid agonist enhances cytotoxic effects in GB7 glioblastoma cancer stem cells. Neurochemistry International 118, 52–60. doi:10.1016/j.neuint.2018.04.010

Dallanoce, C., Conti, P., De Amici, M., De Micheli, C., Barocelli, E., Chiavarini, M., Ballabeni, V., Bertoni, S., Impicciatore, M., 1999. Synthesis and functional characterization of novel derivatives related to oxotremorine and oxotremorine-M. Bioorganic & Medicinal Chemistry 7, 1539–1547. doi:10.1039/C7MD00149E

De Amici, M., Dallanoce, C., Holzgrabe, U., Tränkle, C., Mohr, K., 2010. Allosteric ligands for G protein-coupled receptors: a novel strategy with attractive therapeutic opportunities. Med Res Rev 30, 463–549. doi:10.1002/med.20166

De Angelis, F., Tata, A.M., 2016. Analgesic Effects Mediated by Muscarinic Receptors: Mechanisms and Pharmacological Approaches. CNSAMC 16, 218–226. doi:10.2174/1871524916666160302103033

De Angelis, F., Marinelli, S., Fioretti, B., Catacuzzeno, L., Franciolini, F., Pavone, F., Tata, A.M., 2014. M2 receptors exert analgesic action on DRG sensory neurons by negatively modulating VR1 activity. J. Cell. Physiol. 229, 783–790. doi:10.1002/jcp.24499

De Min, A., Matera, C., Bock, A., Holze, J., Kloeckner, J., Muth, M., Traenkle, C., De Amici, M., Kenakin, T., Holzgrabe, U., Dallanoce, C., Kostenis, E., Mohr, K., Schrage, R., 2017. A new molecular mechanism to engineer protean agonism at a G protein-coupled receptor. Molecular Pharmacology 91, 348–356. doi:10.1124/mol.116.107276

Decker, M., 2017. Design of Hybrid Molecules for Drug Development. Elsevier.

Disingrini, T., Muth, M., Dallanoce, C., Barocelli, E., Bertoni, S., Kellershohn, K., Mohr, K., De Amici, M., Holzgrabe, U., 2006. Design, synthesis, and action of oxotremorine-related hybrid-type allosteric modulators of muscarinic acetylcholine receptors. J. Med. Chem. 49, 366–372. doi:10.1021/jm050769s

Eddy, N.B., Leimbach, D., 1953. Synthetic analgesics. II. Dithienylbutenyl- and dithienylbutylamines. J. Pharmacol. Exp. Ther. 107, 385–393.

Eisenach, J.C., 1999. Muscarinic-mediated analgesia. Life Sci. 64, 549–554. doi:10.1016/s0024-3205(98)00600-6

Fiorino, D.F., Garcia-Guzman, M., 2012. Muscarinic pain pharmacology: realizing the promise of novel analgesics by overcoming old challenges., in: Handbook of Experimental Pharmacology. Springer Berlin Heidelberg, Berlin, Heidelberg, pp. 191–221. doi:10.1007/978-3-642-23274-9_9

Furchgott, R.F., Bursztyn, P., 1967. Comparison of dissociation constants and of relative efficacies of selected agonists acting on parasympathetic receptors. Ann NY Acad Sci 144, 882–899. doi:10.1111/j.1749-6632.1967.tb53817.x

Gould, T.D., Dao, D.T., Kovacsics, C.E., 2009. Mood and anxiety related phenotypes in mice: characterization using behavioral tests 42. doi:10.1007/978-1-60761-303-9

Hayashida, K.-I., Bynum, T., Vincler, M., Eisenach, J.C., 2006. Inhibitory M2 muscarinic receptors are upregulated in both axotomized and intact small diameter dorsal root ganglion cells after peripheral nerve injury. Neuroscience 140, 259–268. doi:10.1016/j.neuroscience.2006.02.013

Jones, P.G., Dunlop, J., 2007. Targeting the cholinergic system as a therapeutic strategy for the treatment of pain. Neuropharmacology 53, 197–206. doi:10.1016/j.neuropharm.2007.04.002

King, N.B., Fraser, V., 2013. Untreated pain, narcotics regulation, and global health ideologies. PLoS Med. 10, e1001411. doi:10.1371/journal.pmed.1001411

Koster, R., Anderson, M., De Beer, E.J., 1959. Acetic acid for analgesic screening. Federation Proceedings 18, 412–417.

Kruse, A.C., Kobilka, B.K., Gautam, D., Sexton, P.M., Christopoulos, A., Wess, J., 2014. Muscarinic acetylcholine receptors: novel opportunities for drug development. Nat Rev Drug Discov 13, 549–560. doi:10.1038/nrd4295

Matera, C., Bono, F., Pelucchi, S., Collo, G., Bontempi, L., Gotti, C., Zoli, M., De Amici, M., Missale, C., Fiorentini, C., Dallanoce, C., 2019. The novel hybrid agonist HyNDA-1 targets the D3R-nAChR heteromeric complex in dopaminergic neurons. Biochem. Pharmacol. 163, 154–168. doi:10.1016/j.bcp.2019.02.019

Matera, C., Flammini, L., Quadri, M., Vivo, V., Ballabeni, V., Holzgrabe, U., Mohr, K., De Amici, M., Barocelli, E., Bertoni, S., Dallanoce, C., 2014. Bis(ammonio)alkane-type agonists of muscarinic acetylcholine receptors: Synthesis, in vitro functional characterization, and in vivo evaluation of their analgesic activity. Eur J Med Chem 75, 222–232. doi:10.1016/j.ejmech.2014.01.032

Matera, C., Pucci, L., Fiorentini, C., Fucile, S., Missale, C., Grazioso, G., Clementi, F., Zoli, M., De Amici, M., Gotti, C., Dallanoce, C., 2015. Bifunctional compounds targeting both D2and non-α7 nACh receptors: Design, synthesis and pharmacological characterization. Eur J Med Chem 101, 367–383. doi:10.1016/j.ejmech.2015.06.039

Matera, C., Tata, A.M., 2014. Pharmacological approaches to targeting muscarinic acetylcholine receptors. Recent Patents on CNS Drug Discovery 9, 85–100. doi:10.2174/1574889809666141120131238

Messerer, R., Dallanoce, C., Matera, C., Wehle, S., Flammini, L., Chirinda, B., Bock, A., Irmen, M., Tränkle, C., Barocelli, E., Decker, M., Sotriffer, C., De Amici, M., Holzgrabe, U., 2017. Novel bipharmacophoric inhibitors of the cholinesterases with affinity to the muscarinic receptors M1 and M2. MedChemComm 8, 1346–1359. doi:10.1039/c7md00149e

Mohr, K., Tränkle, C., Kostenis, E., Barocelli, E., De Amici, M., Holzgrabe, U., 2010. Rational design of dualsteric GPCR ligands: quests and promise. Br. J. Pharmacol. 159, 997–1008. doi:10.1111/j.1476-5381.2009.00601.x

Pan, H.-L., Wu, Z.-Z., Zhou, H.-Y., Chen, S.-R., Zhang, H.-M., Li, D.-P., 2008. Modulation of pain transmission by G-protein-coupled receptors. Pharmacol. Ther. 117, 141–161. doi:10.1016/j.pharmthera.2007.09.003

Riefolo, F., Matera, C., Garrido-Charles, A., Gomila, A.M.J., Sortino, R., Agnetta, L., Claro, E., Masgrau, R., Holzgrabe, U., Batlle, M., Decker, M., Guasch, E., Gorostiza, P., 2019. Optical Control of Cardiac Function with a Photoswitchable Muscarinic Agonist. J. Am. Chem. Soc. 141, 7628–7636. doi:10.1021/jacs.9b03505

Sánchez, C., Meier, E., 1993. Central and peripheral mediation of hypothermia, tremor and salivation induced by muscarinic agonists in mice. Pharmacol. Toxicol. 72, 262–267. doi:10.1111/j.1600-0773.1993.tb01647.x

Tata, A.M., 2008. Muscarinic Acetylcholine Receptors: New Potential Therapeutic Targets in Antinociception and in Cancer Therapy. Recent Patents on CNS Drug Discovery 3, 94–103. doi:10.2174/157488908784534621

Viegas-Junior, C., Danuello, A., da Silva Bolzani, V., Barreiro, E.J., Fraga, C.A.M., 2007. Molecular hybridization: a useful tool in the design of new drug prototypes. Curr. Med. Chem. 14, 1829–1852. doi:10.2174/092986707781058805

Woolfe, G., MacDonald, A.D., 1944. The evaluation of the analgesic action of pethidine hydrochloride (Demerol). J. Pharmacol. Exp. Ther. 80, 300–307. doi:10.1002/(ISSN)2052-1707

